# Spatial orientation based on multiple visual cues in monarch butterflies

**DOI:** 10.1101/2020.02.13.947739

**Authors:** Myriam Franzke, Christian Kraus, David Dreyer, Keram Pfeiffer, M. Jerome Beetz, Anna L. Stöckl, James J. Foster, Eric J. Warrant, Basil el Jundi

## Abstract

Monarch butterflies (*Danaus plexippus*) are prominent for their annual long-distance migration from North America to its overwintering area in Central Mexico. To find their way on this long journey, they use a sun compass as their main orientation reference but will also adjust their migratory direction with respect to mountain ranges. This indicates that the migratory butterflies also attend to the panorama to guide their travels. Here we studied if non-migrating butterflies - that stay in a more restricted area to feed and breed - also use a similar compass system to guide their flights. Performing behavioral experiments on tethered flying butterflies in an indoor LED flight simulator, we found that the monarchs fly along straight tracks with respect to a simulated sun. When a panoramic skyline was presented as the only orientation cue, the butterflies maintained their flight direction only during short sequences suggesting that they potentially use it for flight stabilization. We further found that when we presented the two cues together, the butterflies register both cues in their compass. Taken together, we here show that non-migrating monarch butterflies can combine multiple visual cues for robust orientation, an ability that may also aid them during their migration.

**Summary:** Non-migrating butterflies keep directed courses when viewing a simulated sun or panoramic scene. This suggest that they orient based on multiple visual cues independent of their migratory context.

## Introduction

Despite their tiny brains, insects exhibit incredible orientation behaviors that range from simple compass orientation (Byrne et al., 2003; el Jundi et al., 2019), to more complex behaviors such as path integration (Collett and Collett, 2000; Heinze et al., 2018) or long-distance migration (Dreyer et al., 2018b; Merlin and Liedvogel, 2019; Warrant et al., 2016). One prominent model organism for the study of spatial orientation in the context of migration is the monarch butterfly (*Danaus plexippus*) (Reppert and de Roode, 2018; Reppert et al., 2016). These colorful butterflies migrate every year over more than 4,000 km from North America and Canada to their overwintering habitat in Central Mexico. To find their route, they rely on celestial compass cues, such as the sun and polarized light (Mouritsen and Frost, 2002; Froy et al., 2003; Reppert et al., 2004; Reppert, 2006; Heinze and Reppert, 2011), with the sun being their main orientation reference during migration (Stalleicken et al., 2005). In addition, they compensate their sun compass based on time-of-day information from circadian clocks in the brain (Sauman et al., 2005) and/or the antennae (Guerra et al., 2012; Merlin et al., 2009; Merlin et al., 2011) to keep a constant southerly migratory direction over the entire course of a day (Froy et al., 2003; Mouritsen and Frost, 2002). Besides these cues, observations of the butterflies’ heading directions indicate that they additionally rely on terrestrial cues to adjust their migratory direction (Calvert, 2001). While it is still unclear whether the butterflies use terrestrial cues in combination with skylight cues, it is known that migrating Bogong moths constantly integrate visual landmarks with the Earth’s magnetic field to maintain a directed course (Dreyer et al., 2018a).

To obtain a robust orientation compass, it is well established that many insects use a combination of visual cues from their environment. Ants combine skylight (Lebhardt and Ronacher, 2015; Wehner, 1997) and terrestrial cues, such as the panoramic skyline (Collett and Collett, 2002; Durier et al., 2003; Graham and Cheng, 2009a; Judd and Collett, 1998), to define the desired homeward direction. Integration of multiple visual cues does not require a certain context (migration or homing) but is a common strategy of insects to keep track of their heading with respect to their environment, irrespective of their behavioral state. Neurobiological studies in flying fruit flies showed that the insect’s internal compass encodes the entire visual scene in a highly flexible manner (Fisher et al., 2019; Kim et al., 2019). This highly dynamic coding of visual cues allows an insect to constantly integrate multiple cues, such as a panoramic scenery, in its compass and to set it in relation to the sun’s position. A very similar internal compass network not only steers migration to Mexico in monarch butterflies (Heinze and Reppert, 2011; Heinze et al., 2013) but also likely guides the animals through their environment in the non-migrating phase. Whether the compass of the non-migrating butterfly also uses skylight cues as an orientation reference and whether it integrates all available visual orientation cues from its environment has not yet been explored.

In this study, we investigated how non-migrating monarch butterflies use single visual cues (simulated sun and panoramic skyline) and a combination of these cues for orientation. We presented the cues to the butterflies while the animals were tethered at the center of an LED flight simulator. Interestingly, although these butterflies were not in their migratory state, we found that the butterflies were able to keep a constant heading direction for the entire flight sequence with respect to a simulated sun. When we presented a panoramic skyline to the butterflies, they were also able to keep constant headings with respect to this stimulus, but only did so for short flight periods. Thus, most butterflies seem to use the panoramic skyline for flight stabilization. When the simulated sun and the panoramic skyline were presented together, we found that the butterflies used both cues for orientation. Their directedness dropped if they had only one cue for orientation. Our results show that, irrespective of their migratory or internal state, monarch butterflies maintain a directed heading based on a simulated sun and terrestrial cues. These results will allow us to explore the ability of the butterflies to keep a constant course not only during their migratory phase but also in their non-migratory phase.

## Material and Methods

### Experimental animals

Monarch butterfly (*Danaus plexippus*) pupae were obtained from the butterfly supplier Costa Rica Entomology Supply (butterflyfarm.co.cr). The pupae were reared in an incubator (HPP 110 and HPP 749, Memmert GmbH + Co. KG, Schwabach, Germany) at 25°C and 80% relative humidity under a 12:12 hour light/dark cycle. After eclosion, the adult butterflies were kept inside a flight cage in an incubator (I-30VL, Percival Scientific, Perry, IA, USA) at a 12:12 hour light/dark cycle. The incubator was set to 25° in the light and 23° in the dark phase and to a constant relative humidity of 50%. The butterflies had free access to a feeder containing 15% sucrose solution. In our experiment, we only used adult butterflies 3-12 days after eclosion. For all experiments, we tested a new group of butterflies.

Prior to the experiments, each butterfly’s thorax was cleared of scales and a tungsten stalk (0.508 x 152.4 mm, Science Products GmbH, Hofheim, Germany) was attached to the thorax dorsally using an instant adhesive glue (multi-purpose impact instant contact adhesive, EVO-STIK, Bostik Ltd, Common Road, Stafford, ST16 3EH, UK). Before the butterflies were tethered in the flight simulator, they were kept in a clear plastic container with access to 15% sucrose solution for at least three hours in darkness to allow the glue to harden.

### Flight simulator

All experiments were performed indoors in an LED flight simulator (Fig. 1A). Similar to previous studies (Dreyer et al., 2018a; Dreyer et al., 2018b; Mouritsen and Frost, 2002), the heading directions of individual butterflies were recorded by connecting the tungsten wire to an optical encoder (E4T miniature Optical Kit Encoder, US Digital, Vancouver, WA, USA) at the center of the flight simulator. Body rotations were recorded with a temporal resolution of 200 ms, an angular resolution of 3° and sent via a digitizer (USB4 Encoder Data Acquisition USB Device, US Digital, Vancouver, WA, USA) to a computer with the corresponding software (USB1, USB4: US Digital, Vancouver, WA, USA). To present different visual stimuli to the butterflies, the inner surface of the arena was equipped with an array of 2048 RGB LEDs (16*16 APA102C LED Matrix, iPixel LED Light Co.,Ltd, Baoan Shenzhen, China). The color and intensity of all LEDs was controlled by a raspberry pi (Raspberry Pi 3 Model B, Raspberry Pi Foundation, UK) and a custom written python script.

**Fig. 1.**
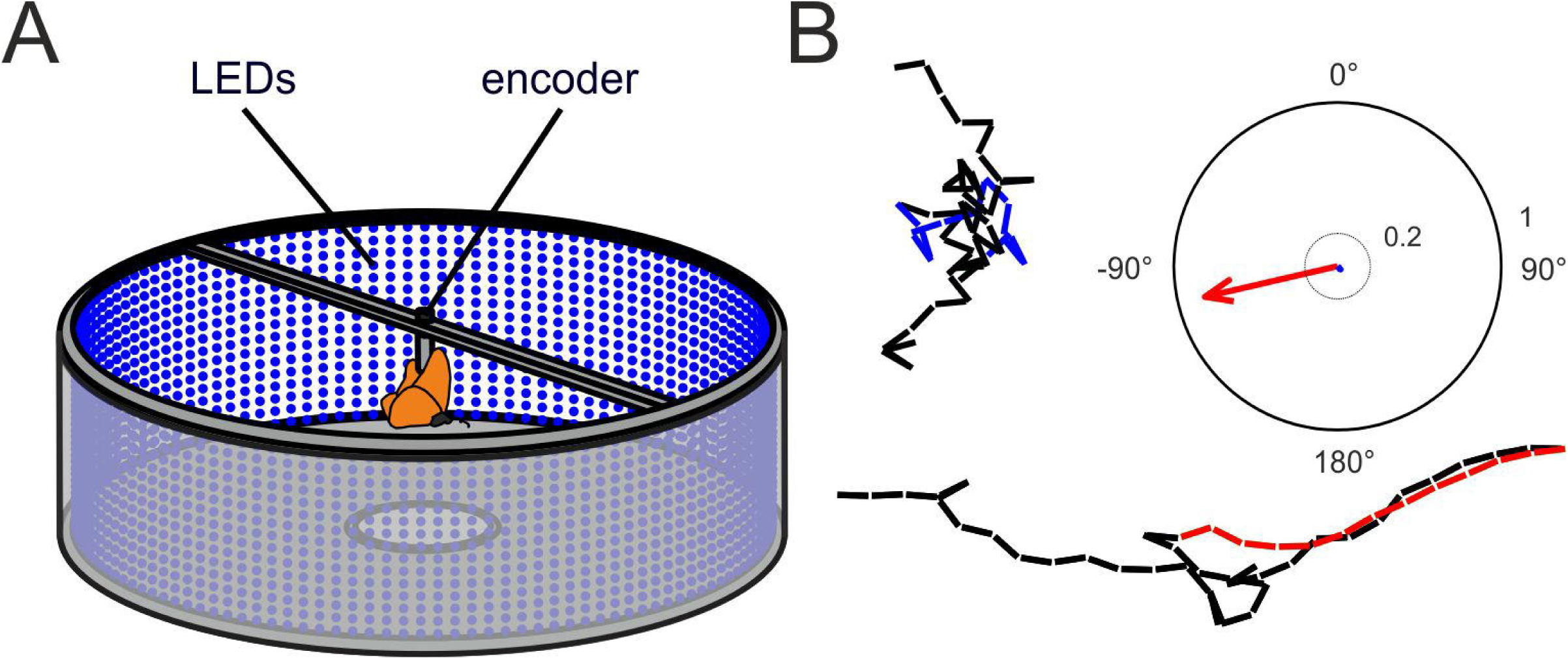
The orientation of tethered monarch butterflies in an LED flight simulator. (***A***) Schematic illustration of a monarch butterfly tethered at the center of the LED flight simulator. The inner surface of the arena is equipped with 2048 RGB-LEDs. While presenting visual stimuli to the butterflies, their heading directions were monitored using an optical encoder. (***B***) Virtual eight-minute flight tracks of a disoriented (upper, left trajectory) and a well-oriented (lower trajectory) butterfly (bin size: 10 s). The red and blue sector of the trajectories indicate a two minutes phase. The red and blue vectors in the circular plot (right) indicate the mean heading direction and vector length *r* of the corresponding phases shown in the fight tracks. The length of the vectors can vary between zero (disoriented) and one (perfectly oriented). The inner dashed circle indicates a vector length of 0.2 and the perimeter of the plot a vector length of 1.

### The sun compass in butterflies

To simulate the sun, one LED, at an elevation between 5-10°, was set to green light (emission peak at approximately 516 nm; intensity of ∼5.2 x 10^12^ photons/cm^2^/s, measured at the center of the arena) while the remaining LEDs of the arena were set to blue light (emission peak at about 458 nm; intensity of ∼4.61 x 10^10^ photons/cm^2^/s/LED, measured at the center of the arena; experiment: *green sun*). Individual butterflies were tethered at the center of the arena and their headings were recorded for eight minutes (Fig. 2B). The position of the stimulus was switched by 180° every two minutes, to ensure that the animals relied on the stimulus presented for orientation. The start position of the sun stimulus was pseudorandomized. Thus, half of the butterflies experienced the sun stimulus at 0° first (0°/180°/0°/180°), while for the other half of the butterflies the sun stimulus was set at 180° first (180°/0°/180°/0°).

**Fig. 2.**
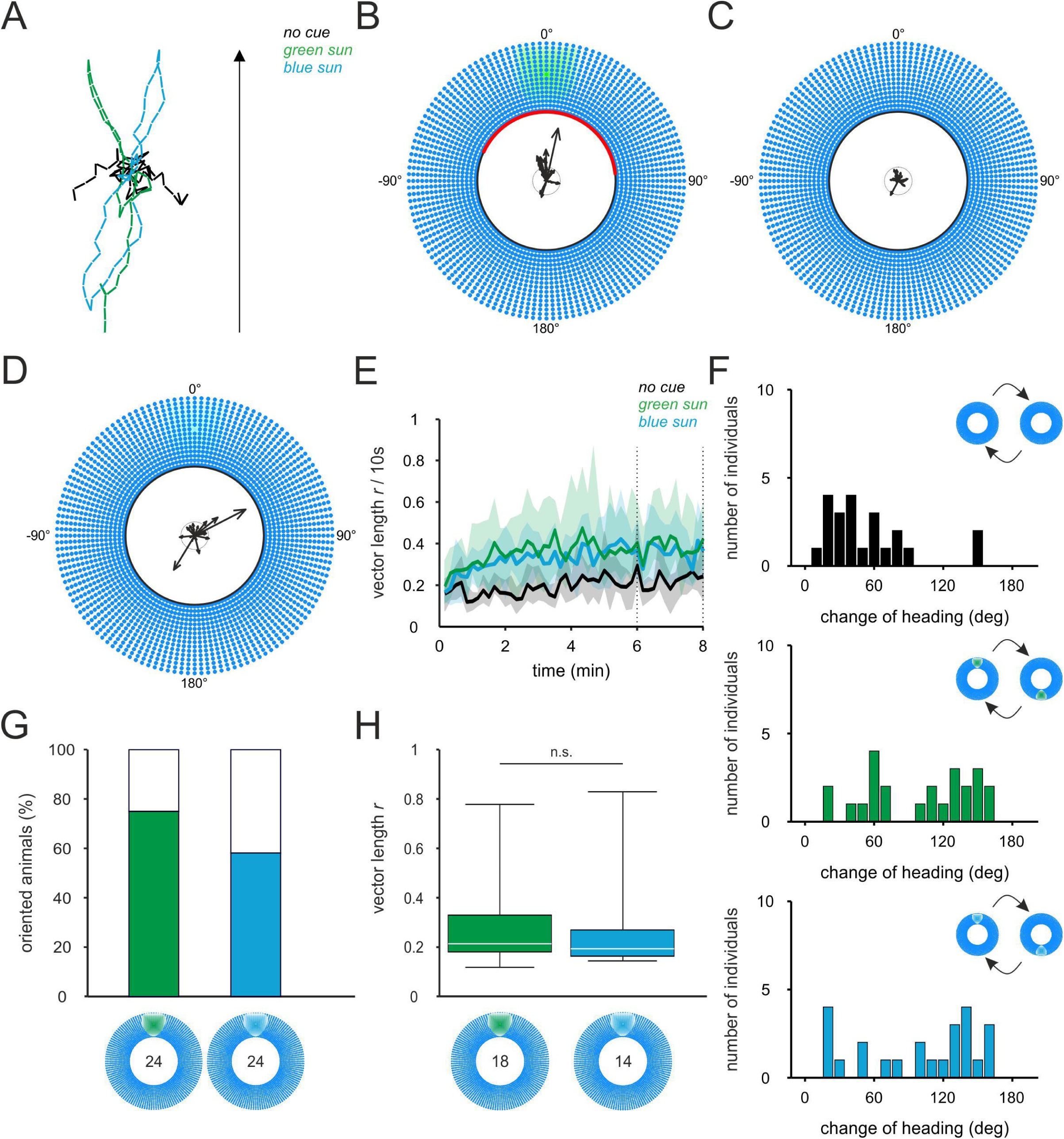
The sun compass in monarch butterflies. ***(A***) Flight trajectories of individual butterflies that viewed a bright *green sun* (green), a bright *blue sun* (blue) or *no cue* (black) as orientation reference. When the sun stimulus was relocated by 180°, the butterflies followed the change of the stimulus’s position. Black arrow indicates the position of the sun stimulus or a specific point in the control scenery in the beginning of the experiment. (***B-D***) Orientation of butterflies with respect to a *green sun* (B; N = 24), without any compass cues (C; N = 22) or a *blue sun* (D; N = 24). The mean vector for every butterfly was calculated over a two-minute phase. The inner circle of the plots indicates *r* = 0.2. The red sector shows the circular standard deviation (SD) of the animals’ significant group orientation. (***E***) The mean vector length *r* (bin size: 10 s) over the entire experiments shows that the butterflies were better oriented with respect to a sun stimulus (green and blue curves) compared to the control without any directional information (black curve). Shaded areas indicate the 25-75% quantile. The vertical dashed lines indicate the two-minute sector that was used to present the heading direction in B-D. (***F***) Histogram of heading changes after a 180° relocation of the stimulus (bin size: 5°) for the *no cue* (upper plot), the *green sun* (middle plot) and *blue sun* (lower plot) experiment. (***G***) The number of “oriented” butterflies was calculated by analyzing which animals showed a vector length *r* > 0.1169 (which is mean plus the 95% confidence interval of the no cue experiment, i.e. the data shown in C). (***H***) The mean vector length did not differ significantly between the experiments with the green and blue sun stimulus (p = 0.43, χ^2^ = 0.64; Kruskal-Wallis-Test). White horizontal lines indicate the median vector length. The boxes show the interquartile range and whiskers extend to the 2.5^th^ and 97.5^th^ percentile. n.s. indicates a difference that is not significant: p > 0.05.

To understand which features, the spectral or intensity information, of the sun stimulus butterflies used, we performed an additional experiment [again over eight minutes (Fig. 2D)] in which we excluded the spectral information from the sun stimulus. Thus, the animals’ behavior was tested by performing the same experiment as with the *green sun* but with a blue LED that had the same spectrum as the remaining blue LEDs of the arena but was much brighter (∼5.2 x 10^12^ photons/cm^2^/s, measured at the center of the arena; experiment: *blue sun*).

To exclude the possibility that butterflies used any additional cues in the experimental setup or room, a negative control experiment was performed for eight minutes with all LEDs set to the same blue wavelength and intensity (experiment: *no cue;* Fig. 2C).

### The use of a panoramic skyline

To investigate how butterflies orient with respect to a terrestrial cue, we presented the animals with a panoramic skyline with smaller and higher profiles (Fig. 3), while the background above the horizon was set to blue light (emission peak at 458 nm) and an intensity of ∼4.61 x 10^10^ photons/cm^2^/s/LED *panorama*). The butterflies’ headings were recorded for eight minutes while the position of the stimulus switched by 180° every two minutes. In a control experiment, the profile of the panorama was removed by switching off LEDs below an elevation of ∼0°, which resulted in a flat horizon (*no profile*). To gain a deeper understanding of how the butterflies used the presented simulated panoramic skyline, compass orientation vs. flight stabilization, we performed an experiment with a stationary grating of vertical stripes in blue (emission peak at approximately 458 nm; three columns of LEDs per stripe, spatial frequency of ∼0.06 cycles/degree; experiment: *grating*) and black. Each blue LED had an intensity of ∼4.61 x 10^10^ photons/cm^2^/s. The flight performance of the butterflies was recorded for four minutes.

**Fig. 3.**
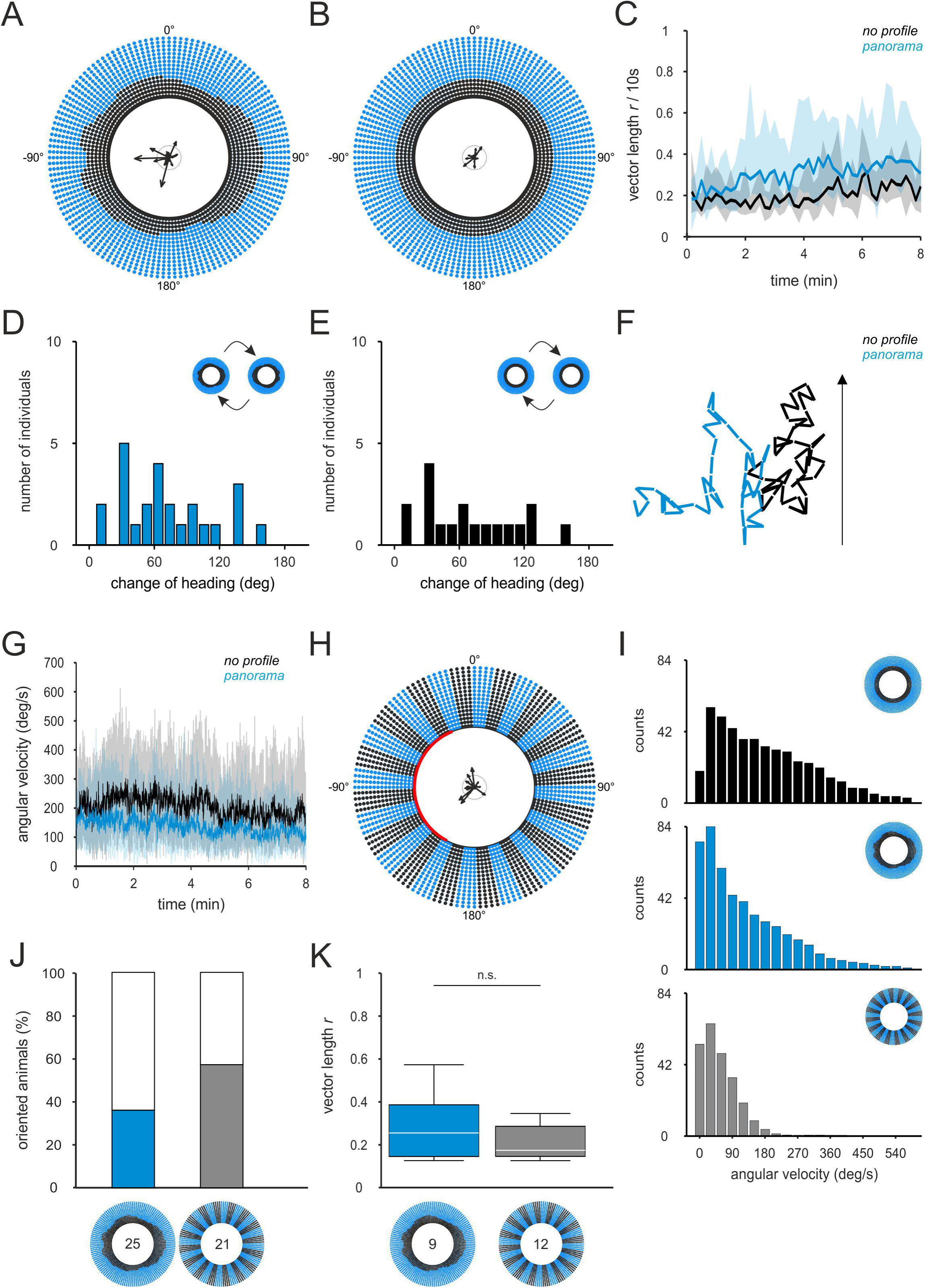
Using a panoramic skyline for a directed flight behavior. (***A-B***) Orientation of monarch butterflies with respect to a panoramic skyline with (A; N = 25) or without (B; N = 18) a profile. Dashed inner circle shows an *r* = 0.2. (***C***) Over the entire experiment, the mean vector length *r* (bin size: 10s) was always higher when the profile of the panoramic skyline was visible to the butterflies. Shaded areas indicate the 25-75% quantile. (***D, E***) Histogram of heading changes after a 180° relocation of the stimulus (bin size: 5°) for the panorama experiments with profile (D) and without profile (E). (***F***) Exemplary flight trajectories of one animal that flew with respect to a panoramic skyline with profile (blue trajectory) and one that oriented to a panoramic skyline without profile (black trajectory). Black arrow indicates the position of a specific point in the visual scenery in the beginning of the experiment. (***G***) The angular velocity of the animals over the eight minutes flight. The angular velocity decreased when the profile of the panoramic skyline was visible (blue curve) compared to when no profile was available (black curve). The shaded area indicates the 25-75% quantile. (***H***) The orientation of butterflies with respect to a grating pattern (N = 21). Dashed inner circle indicates a vector length of 0.2. The red sector shows the circular SD of the animal’s group orientation. (***I***) The frequency of observed angular velocities (window size of each bin: 30°) when the butterflies had a panoramic skyline without profile (upper plot, same data as G), the panorama with profile (middle plot, same data as G) or a grating pattern as orientation reference. (***J***) The number of “oriented” butterflies was defined as r > 0.1194 (which is the 95% confidence interval of the panorama experiments without profile, i.e. the data shown in B, C). (***K***) The vector length of the “oriented” animals did not differ significantly between the experiments with the skyline panorama and the grating pattern as visual stimulus (p = 0.48, χ^2^ = 0.51; Kruskal-Wallis-Test). White horizontal lines indicate the median vector length. The boxes show the interquartile range and whiskers extend to the 2.5^th^ and 97.5^th^ percentile. n. s. indicates no significant difference (p > 0.05)

### Combination of terrestrial and sun compass information

To answer the question of whether monarch butterflies combine different visual cues to increase their flight accuracy, we presented the panoramic skyline together with the bright green sun stimulus (experiment: *panorama and sun*, Fig. 4). Each butterfly’s orientation performance was recorded for eight minutes and the position of the scenery was switched by 180° every two minutes. In a control experiment where the profile of the panorama lacked any bumps (i.e. was a flat horizon), the panoramic features were excluded but the sun was available (experiment: *sun and no profile*).

**Fig. 4.**
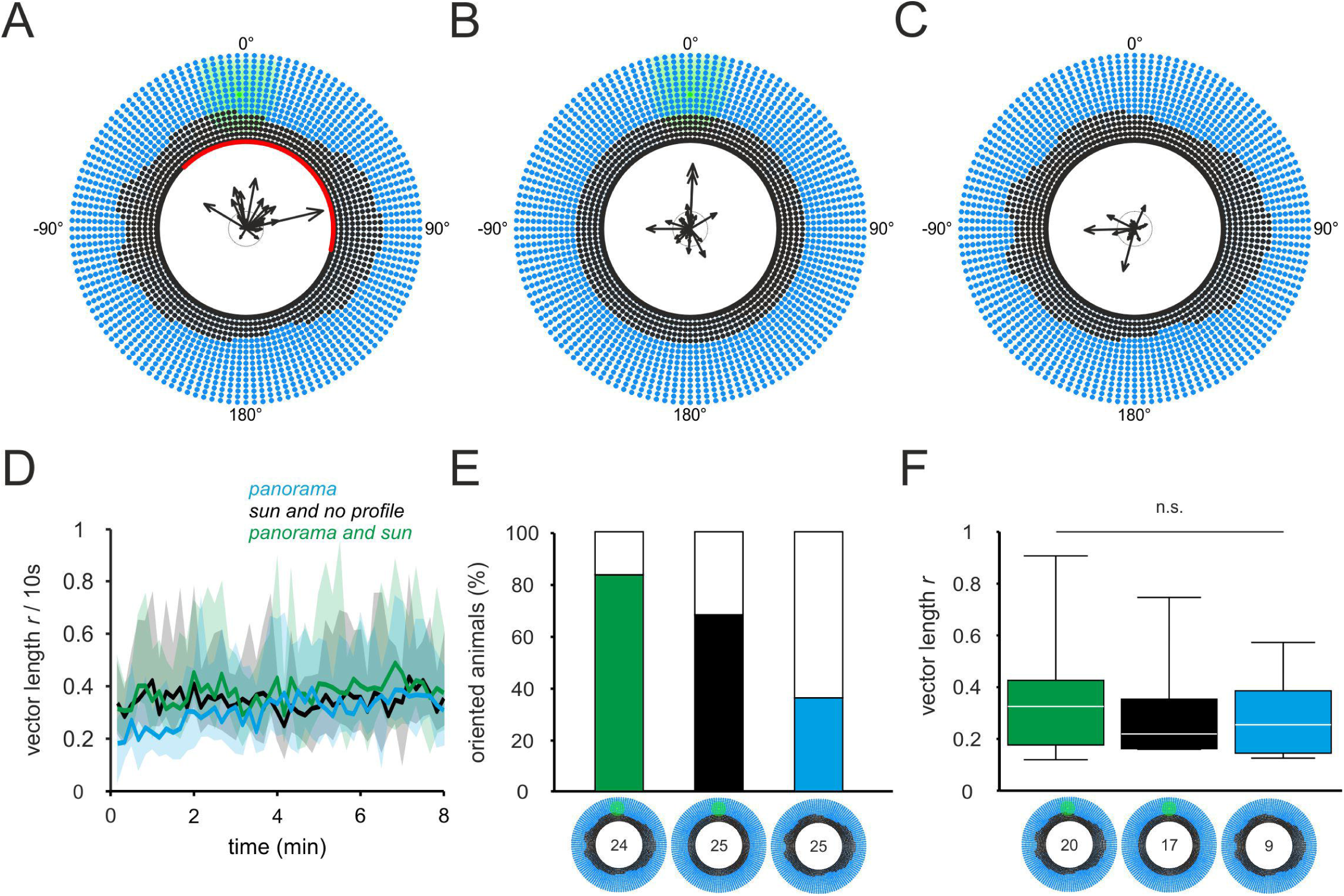
Combining different visual cues for orientation. (***A-C***) The Orientation of butterflies with respect to the panoramic skyline with (A; N = 24) or without (B; N = 25) the profile combined with a green sun or the panoramic skyline alone (C; N = 25, same data as shown in Fig. 3A). Dashed inner circle of the circular plots indicate an *r* value of 0.2. The red sector in A indicates the SD. (***D***) Over an eight-minute flight, the mean vector length *r* was relatively high irrespective of whether only one of the cues, the panorama (blue curve) or the sun (black curve) or both panorama and sun (green curve) were available. Shaded areas show the 25-75% quantile (***E***) The number of animals with a vector length over *r* > 0.1194 (which is the 95% confidence interval of the control experiment *no profile*). (***F***) The orientation performance of the “oriented” animals did not differ significantly between the experiments with one of the cues (blue box plot = panorama only; black box plot = sun only) or both cues available (green box plot) (p = 0.61; χ^2^ = 0.99; Kruskal-Wallis-Test;). White horizontal lines represent the median vector length *r*. The boxes show the interquartile range and whiskers extend to the 2.5^th^ and 97.5^th^ percentile. n.s. indicates no significant difference (p > 0.05).

In an additional experiment, we investigated how the disappearance of a visual cue affects the butterflies’ orientation performance (Fig. 5). We first allowed the butterflies to acclimate to the experimental conditions for two minutes (with the green sun and panorama available) as we noticed in the sun and panorama experiments (Figs. 2E, 3C) that the orientation abilities of butterflies significantly changed over the first two minutes. In the subsequent 30 seconds, the butterflies were again presented with the combination of the panoramic skyline and the green sun stimulus (*combination*). For the next 30 seconds, we excluded one of the stimuli (we either removed the sun stimulus or removed the profile of the panorama (*single cue*). Half of the animals were first tested without a panorama (but with the sun), while half of the butterflies first experienced the panorama (without the sun). All butterflies experienced both stimuli again for an additional 30 seconds (*combination*) before the other stimulus that was present in phase 2 (either the simulated sun or the panorama) was removed (*single cue*) for further 30 seconds. The order of the stimulus presentation (both cues/simulated sun, both cues/panorama) was pseudorandomized.

**Fig. 5.**
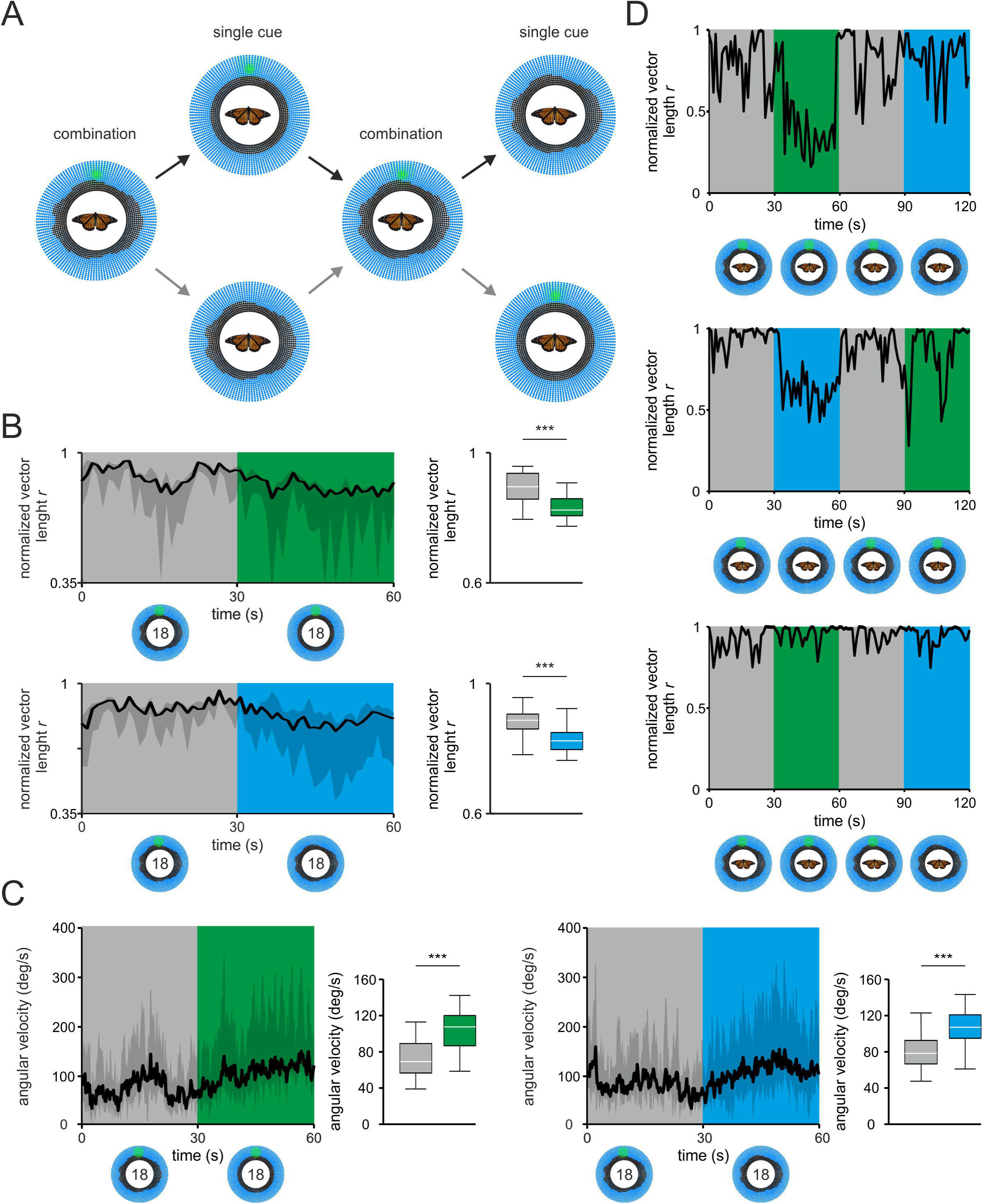
Monarch butterflies can use a combination of different cues for orientation. (***A***) Schematic illustration of the experimental procedure. We tested the use of different cues by presenting both cues (panorama and simulated sun, phase 1) to the butterflies and subsequently removed one of the cues (phase 2), either the profile of the panorama (black arrow) or the sun (grey arrow). In the third phase, we presented both cues to the butterflies before one cue was withheld in the fourth phase. Each phase lasted for 30 seconds. (***B***) On average the butterflies showed a significant decrease in their directedness when either of the cues was excluded. Left plot: normalized vector length (per 1 s) when the panorama (upper panel) or the simulated sun (lower panel) was excluded after 30 s of flight. Grey shaded areas in the left plots represent the 25%-75% quartile. Right plot: Box plots of the averaged normalized vector length (same data as in left plot). The normalized vector length dropped significantly when the panoramic skyline (upper plot, p < 0.001, Z = 4.33; Wilcoxon signed-rank test) or the green sun (lower plot, p < 0.001, Z =3.71; Wilcoxon signed-rank test) disappeared. *** indicates a significant difference of p < 0.001. (***C***) The angular velocity increased when the panorama (left panel) or the simulated sun (lower panel) disappeared (same data as in C). The velocity increased significantly when the green sun (left box plot, p < 0.001, Z = −9.40; Wilcoxon signed-rank test) or the panorama (right box plot, p < 0.001, Z = − 7.90; Wilcoxon signed-rank test) was suddenly the only available orientation reference. Grey shaded areas in the left plots represent the 25%-75% quartile. Box plots: white horizontal lines represent the median vector length. The boxes indicate the interquartile range and whiskers extend to the 2.5^th^ and 97.5^th^ percentile. *** indicates a significant difference of p < 0.001. (***D***) The normalized vector length (per 1 sec.) of oriented, individual butterflies is shown over the 2 minutes experiment. Individual butterflies used different cues for orientation. Some butterflies relied on one of the cues, panorama (upper panel) or the simulated sun (middle panel), and their performances decreased when this cue was excluded. A few butterflies used both cues and showed a high orientation throughout the entire experiment (lower panel).

## Data analysis

All data were analyzed in the software MATLAB (Version R2017b, MathWorks, Natick, MA, USA) using the CirCStat toolbox (Berens, 2009). The experiments that took eight minutes (*green sun, blue sun, no cue, panorama, no profile, panorama and sun, no profile and sun*; Figs. 2-4) were divided into four phases of equal length, and all butterflies that stopped more than four times during the experiments were excluded from the analysis. The *grating* experiment (Fig. 3) was split into two phases (two minutes each). Because this experiment lasted for only four minutes, butterflies that stopped flying more than two times were excluded from the analysis. This exclusion criterion was also used for the combination experiment (Fig. 5). Depending on the experiment, the data were divided into either two-minute (Figs. 2-4) or 30 second (Fig. 5) phases.

To present the data with respect to the stimulus position, all heading directions were shifted in such a way that the simulated sun or a specific point of the panorama stimulus was positioned at 0°. For each butterfly we calculated the flight trajectory (e.g. in Fig. 1B), and the mean vector within each ten-second bin and within a phase (two-minute bins) (Figs. 2-4). The mean direction *µ* of each butterfly within a phase was calculated (Fig. 1B). To obtain the animal’s performance on a finer scale in the *combination* experiment (Fig. 5), we calculated the vector length *r* within a window size of one second. To avoid any misinterpretation of these *r* values (they are higher than the *r* values over 10 sec. or 2 min.), we normalized all *r* values to the highest *r* value obtained in each flight. To further analyze the butterflies’ performance in our flight simulators, we calculated the angular velocities of the butterflies. A highly oriented animal shows low angular velocities, usually caused by slow swinging around the heading direction. Highly disoriented animals exhibit high angular velocities, often caused by rapid rotation. The angular velocity of individual butterflies was defined by calculating the absolute angular difference between two consecutive headings (Figs. 3, 5). To further test whether the butterflies followed a relocation of a stimulus, their change of heading was calculated by measuring the angular differences in the mean direction between two consecutive phases (Figs. 2F, 3D, 3E). As each individual experienced three stimulus relocations over the 8-minutes flight (after 2, 4, and 6 minutes), we calculated the mean change of heading over three stimulus-relocations in each animal.

## Statistics

During our experiments, we noticed that many butterflies exhibited very poor performance in the first two minutes as compared to the remaining six minutes (e.g. *green sun* and *blue sun*, Fig. 2E). Depending on the experiment, the butterfly’s ability to keep a constant course continued to improve considerably over the first phases. To ensure that we compare the butterflies at a phase when they had adjusted to the experimental situation, we focused on the last phase of each experiment for the statistical evaluation. A possible bias of the heading directions towards a certain direction within this phase was tested with the non-parametric Moore’s Modified Rayleigh test (Moore, 1980). Furthermore, some butterflies performed poorly, for example failing to follow the change of the stimulus’ position, most likely because they were unmotivated to use the visual scene for orientation. To compare the performance of the butterflies, we first calculated the mean *r* within the last two-minute phase of the control experiments plus the 95% confidence interval (*no cue*: *r* = 0.1169, *no profile*: *r* = 0.1194). All animals that showed a lower directedness than these *r* values were excluded from the comparison. The performance of the butterflies was statistically compared using a Kruskal-Wallis-Test for samples of different groups or using the Wilcoxon signed-rank test for comparison within the same group of butterflies (e.g. Fig. 5). The Mardia-Watson-Wheeler test was used to compare the heading directions of different butterfly groups.

## Results

### The sun compass in butterflies

To study the orientation of monarch butterflies with respect to a simulated sun, we recorded the flight performance while the animals were tethered at the center of the LED flight simulator (Fig. 1A) and were presented with a green, bright light spot against a blue background as their only orientation reference. Many monarch butterflies, even though they were outside of their migratory phase, kept a constant heading direction with respect to this stimulus. When the stimulus’s position was turned by 180°, these butterflies changed their heading accordingly (Fig. 2A, green trajectory). On average, the butterflies chose headings towards the simulated sun (p = 0.002, R = 1.46; *µ* = 9° with respect to the simulated sun; non-parametric Moore’s Modified Rayleigh test; *green sun*; N = 24; Fig. 2B). Next, we switched the green stimulus LED to blue, so it was indistinguishable from all other LEDs in the arena. Unsurprisingly, the *r* values, which describe the orientation precision of each butterfly across the two-minute phase, were significantly lower in the absence of the simulated sun (*no cue*; N = 22; Fig. 2C) than when the sun stimulus was available (p = 0.001, χ^2^ = 10.59; Kruskal-Wallis-Test; Fig. 2B). This was also evident when we analyzed the flight directedness on a much finer temporal scale (Fig. 2E): the vector length *r*/10 s increased from on average 0.30 ± 0.16 (mean ± SD) over the first two minutes when the animals viewed the simulated sun and remained stable at a vector length of about 0.39 ± 0.20 for the subsequent six minutes of flight (*green sun*; Fig. 2E). In contrast, the vector length remained relatively low (at 0.20 ± 0.10) throughout the entire eight-minute flight in the absence of any cue (*no cue*; Fig. 2E). Taken together, the improvement of orientation in the presence of a directional stimulus and the following of the stimulus show that non-migrating butterflies use the sun stimulus in our flight simulator for orientation.

To investigate whether the butterflies relied on the spectral or the brightness component of the sun stimulus for orientation, we presented the simulated sun as a bright, blue spot. Similar to what we observed with the *green sun*, butterflies were able to keep a directed course with respect to the *blue sun* and changed their heading when the stimulus was displaced by 180° (Fig. 2A, blue trajectory). The heading directions of the butterflies in the *blue sun* experiment were uniformly distributed (p = 0.22, R = 0.71; non-parametric Moore’s Modified Rayleigh test; *blue sun*; N = 24; Fig. 2D). The vector length *r*/10 s over the entire flight sequence exhibited a similar time course for the experiments with the *green sun* and *blue sun* (Fig. 2E). Although many butterflies kept a constant course in both experiments, we noticed that only a subpopulation of animals followed the 180° relocation of the stimulus [14 out of 24 (*green sun*) and 15 out of 24 (*blue sun*) showed a change in heading > 90°; Fig. 2F]. The remaining butterflies did not change their heading as expected if they used the presented cues for orientation. To exclude any potential effects due to differences in the butterflies’ behavioral state, we analyzed how many animals exhibited a higher *r* value under conditions with a cue (*green sun* and *blue sun*) compared to the mean [plus 95% confidence interval (CI)] *r* values when no visual cue (*no cue*) was available. 18 out of 24 butterflies (75%) presented with the *green sun* stimulus showed a higher mean vector length *r* than when the cue was absent (*r* > 0.1169), while only 14 out of 24 animals (58%) showed a higher vector length in the *blue sun* experiment (Fig. 2G). The vector length *r* of these “oriented” animals was similar in the *green sun* and *blue sun* experiment (p = 0.43, χ^2^ = 0.64; Kruskal-Wallis-Test; Fig. 2H) and no significant differences in the heading directions between both groups were found (p = 0.06, W = 5.64; Mardia-Watson-Wheeler test). This suggests that a sun stimulus that contains only brightness information elicits a similar ability to keep a constant heading as a stimulus that contains both spectral and brightness information.

### The use of a panoramic skyline

Next, we presented a panoramic skyline to the animals with the panorama’s profile consisting of smaller and higher bumps (Fig. 3A). The animals kept arbitrary headings with respect to the panorama stimulus (p = 0.37, R = 0.58; non-parametric Moore’s Modified Rayleigh test; *panorama*, N = 25; Fig. 3A). When the panorama was flat, i.e. the panorama’s profile was absent, we observed that none of the tested animals kept a constant heading (*no panorama*, N = 18; Fig. 3B). Although the length of mean vector *r* was similar when the panorama exhibited a profile (Fig. 3A) compared to when no profile (Fig. 3B) was available (p = 0.12, χ^2^ = 2.41; Kruskal-Wallis-Test), the vector lengths *r*/10 s over the entire experiment was significantly higher with the profile as compass cue (p < 0.001, χ^2^ = 13.53; Kruskal-Wallis-Test; Fig. 3C). This suggests that the panoramic skyline stimulus improves the butterfly’s ability to maintain a directed course.

We also noticed that the butterflies did not change their headings when the visual scene was turned by 180° (Figs. 3D, 3E). In general, they did not keep a certain heading direction with respect to the panorama stimulus (as in the sun experiments) but rather constantly changed their headings over the course of the experiment (Fig. 3F). This opens up the possibility that the panorama might not be used for compass orientation but for flight stabilization. This is further supported by the observation that the rotational speed of the animals, i.e. the angular velocity, was significantly lower when the panoramic features were available to the animals (p < 0.0001, χ^2^ = 2633.71; Kruskal-Wallis-Test; Fig. 3G). To next study the butterflies’ performance with respect to a visual stimulus that provides a strong rotational optic flow but lacks any directional information, we conducted an additional experiment in which we presented a stationary grating pattern to the butterflies (*grating*; N = 21; Fig. 3H). Interestingly, while the butterflies showed angular velocities up to about 540°/s in both panorama experiments (*panorama* and *no profile*), the animals’ angular velocity did not exceed 180°/s in the *grating* experiment (Fig. 3I). This demonstrates that optic-flow information is perceived by the monarch butterflies in this setup and plays an important role while the animals aim to keep a constant flight direction.

To compare the butterflies’ orientation performance between the *panorama* and *grating* experiment, we first analyzed how many animals exhibited a higher *r* value in these experiments than in the *no profile* (mean *r* + 95% CI) experiment. While only 9 out of 25 animals (36%) showed a higher *r* value in the experiment with the *panorama*, 12 out of 21 (57%) were higher in the *grating pattern* experiment (Fig. 3J). The mean vector length *r* of the “oriented” animals did not differ between both experiments (p = 0.48, χ^2^ = 0.51; Kruskal-Wallis-Test; Fig. 3K). Nevertheless, we noticed that two butterflies exhibited a very high directedness (r > 0.4; Figs. 3A, K) which was never observed in the grating pattern experiments (Figs. 3H, K).

### Combination of terrestrial and sun compass information

To characterize orientation performance in the presence of both terrestrial and celestial cues, we presented the green sun combined with the panoramic skyline including a profile (*panorama and sun*; N = 24; Fig. 4A) or without a profile (*sun and no profile*; N = 25; Fig. 4B). When the profile of the panoramic skyline was added to the scenery (*panorama and sun*) most of the animals kept a specific heading which was centered in a direction between the simulated sun and the center of the highest part of the panorama (p < 0.001, R = 1.63; *µ* = 29° with respect to the simulated sun; non-parametric Moore’s Modified Rayleigh test; Fig. 4A). In contrast, the butterflies chose arbitrary headings in the *sun and no profile* experiment (p = 0.34, R = 0.60; non-parametric Moore’s Modified Rayleigh test; Fig. 4B). This effect was also observed in the experiment without the green sun (*panorama*; Figs. 3A, 4C). The vector length *r*/10 s was stable over the entire flight when both, the sun and panorama, were available [0.38 ± 0.20 (mean ± SD); *panorama and sun*; Fig. 3D] and was significantly higher compared to the condition without the sun [0.30 ± 0.19 (mean ± SD); *panorama*; p = 0.005, χ^2^ = 8.07; Kruskal-Wallis-Test; Fig. 4D)]. However, the directedness did not differ between the *panorama and sun* and when the panorama’s profile was absent [0.35± 0.20 (mean ± SD); *sun and no profile*; p = 0.33, χ^2^ = 0.94; Kruskal-Wallis-Test; Fig. 4D]. To compare the performance of the butterflies, we calculated how many animals exhibited a higher directedness *r* value in the *sun and no profile* and *panorama and sun* experiment compared to the mean r value + 95% CI (*r* = 0.1194) in the *no profile* experiments (Fig. 3B). Irrespective of the panorama, more animals showed higher *r* values as soon as the simulated sun was available (Fig. 4E). The performance of these “oriented” animals did not show any significant differences (p = 0.61; χ^2^ = 0.99; Kruskal-Wallis-Test; Fig. 4F) which suggests that combining different visual cues does not necessarily help to improve the directedness of the butterfly’s flight behavior.

The previous experiment did not allow us to test whether the butterflies registered both visual cues in their compass or if they relied on the simulated sun as their only reference (while ignoring the panoramic skyline). We therefore performed an experiment in which we presented both cues (green sun and panorama; *combination*; Fig. 5A) to the butterflies in a first phase and subsequently withheld one of the cues (*single cue*; Fig. 5A) during a second phase (followed by an additional phase with both cues – *combination* – and a subsequent disappearance of the other cue – *single cue* – see Fig. 5A). 18 of 25 butterflies showed a performance with higher *r* values in the two *combination* phases than in the *no profile* (Fig. 3C) experiment. When analyzing the switch from the two visual stimuli to one cue in these 18 animals, we found that irrespective of which cue we turned off, this led to a significant decrease in the directedness *r* of the butterflies (p < 0.001, Z = 4.33 when the panorama’s profile disappeared; Wilcoxon signed-rank test; Fig. 5B upper panel; p < 0.001, Z = 3.71 when the simulated sun disappeared; Wilcoxon signed-rank test; Fig. 5B lower panel). Associated with this drop in the vector length, the angular velocity increased when only one cue was available (p < 0.001, Z = −9.40 when the panorama’s profile disappeared; Wilcoxon signed-rank test; p < 0.001, Z = −7.90 when the simulated sun disappeared; Wilcoxon signed-rank test; Fig. 5C) further confirming that both cues are being registered by the butterflies. Interestingly, the disappearance of a specific cue had different effects in different animals. In several animals (8 out of 18), we found a drop in the vector length *r* when one of the cues – the simulated sun *or* the panorama – was excluded from the visual scenery (Fig. 5D, upper and middle). In other butterflies (10 out of 18), the disappearance of neither one of the cues had any effect on the directedness (Fig. 5D, lower panel) which indicates that they can dynamically switch from one to another as main orientation reference. Taken together, the data show that monarch butterflies can register multiple visual cues to keep a directed course. However, the relevance of these cues seems to differ in the tested animals.

## Discussion

### The sun compass

Our experiments show that monarch butterflies – even when they are not in their migratory phase – use a green light cue (simulated sun) to keep a constant heading. Thus, keeping track of their heading direction with respect to the sun might not only be an important feature during migration but also to find suitable flower patches in the non-migrating phase. Whether non-migrating butterflies also shift their heading direction in a time-dependent manner to the simulated sun, as the migratory phase does to the real sun (Froy et al., 2003; Mouritsen and Frost, 2002), remains to be investigated.

Our results show that monarch butterflies sometimes prefer to choose a heading towards the simulated green sun (Fig. 1B), while in some experiments they took arbitrary headings (Fig. 4B). The latter is similar to the findings in the fruit fly *Drosophila melanogaster* that also maintains arbitrary headings (Giraldo et al., 2018) and suggest that they are able to perform compass orientation with respect to the sun stimulus. The heading choices towards the green sun in Fig. 2B may potentially result from a reduced ability of the butterflies to detect the sun stimulus in front of the bright blue background. In previous experiments, the sun was presented in front of a dark background (el Jundi et al. 2015b, Giraldo et al. 2018), while in our experiments the illuminated background led to a reduced contrast between the sun stimulus and the background (the sun stimulus was only 2 orders of magnitude brighter than the background). This may results in heading directions where the animals keep the stimulus frontally in their visual fields. This is line with the arbitrary heading choices in Fig. 4B where less blue LEDs were turned on and, thus, a stronger contrast between the sun and blue background was presented to the butterflies. It will now be interesting to test what heading choices the non-migrating butterflies prefer if we study them with respect to the real sun outdoors. But why do the butterflies, even when they are not in their migratory phase, keep constant headings in the flight simulator and what is their behavioral state? Our current interpretation is that the butterflies exhibit an escape response and use the sun stimulus as a reference. An alternative explanation is that the butterflies’ goal, on their search for food, is to disperse into a new niche, a behavior that is well established for butterflies under natural conditions (Felt, 1925; Stevens et al., 2010).

Our experiments show that monarch butterflies use a sun stimulus that contains only brightness information in a very similar way as a stimulus that contains both spectral and brightness information. This suggest that intensity information of the sun can be used by the butterflies to keep a directed course, which is in line with electrophysiological studies that indicate a wavelength-independent neural coding of the sun in the monarch butterfly’s brain (Heinze and Reppert, 2011). In nature, due to a different ratio of shorter (UV light) and longer (green light) wavelengths of light between the solar and anti-solar hemisphere, the direction of the sun can be determined based on a spectral contrast (Coemans et al., 1994; el Jundi et al., 2014). Whether monarch butterflies can use this spectral information, similar to what has been shown for bees (Brines and Gould, 1979; Edrich et al., 1979; Rossel and Wehner, 1984) and dung beetles (el Jundi et al., 2015a; el Jundi et al., 2016) remains to be shown in further experiments. In bees, a green light cue is interpreted as the sun while a UV light cue is treated as a patch of the sky somewhere in the anti-sun direction (Brines and Gould, 1979; Edrich et al., 1979; Rossel and Wehner, 1984). Unfortunately, our current LED stimulus does not allow testing for this in monarch butterflies as the stimulus lacks any UV light, which might be essential for the use of spectral cues in insects. Thus, to fully determine whether monarch butterflies use spectral information for orientation, we aim to additionally present UV light in our flight simulator in the future.

### The use of a panoramic skyline

Monarch butterflies rely on a sun compass (Mouritsen and Frost, 2002; Stalleicken et al., 2005), and potentially also use terrestrial cues to keep a desired heading direction in a similar way as it has been shown in the past for other insects (Cartwright and Collett, 1983; Collett and Land, 1975; Fleischmann et al., 2018; Lehrer and Collett, 1994). Ants and wasps are well-known to use the panoramic skyline as an orientation reference during homing (Graham and Cheng, 2009a, 2009b; Philippides et al., 2011; Reid et al., 2011; Narendra et al., 2013; Narendra and Ramirez-Esquivel, 2017; Stürzl et al. 2016). Calvert (2001) observed a change in the butterfly’s migratory direction as soon as they reached the mountains of the Sierra Madre Orientation. The author suggested that the butterflies might use the beneficial wind conditions generated by the mountain ranges to migrate toward Mexico (Calvert, 2001). Similarly, it was suggested in another study that the Rocky Mountains act as a physical barrier and funnel the butterflies towards Mexico (Mouritsen et al., 2013). In these cases, the animal’s compass can obtain a higher robustness for the maintenance of the migratory direction by combining and matching the sun’s position with the terrestrial scenery. It will therefore be very interesting to test if terrestrial cues play a major role in the context of migration.

Terrestrial cues might especially be relevant if the animals orient in their natural habitat in their non-migratory phase, e.g. during foraging. We therefore presented a dark silhouette of a panoramic skyline to the butterflies as findings in ants suggest that the contrast of objects against the sky is important for the animals’ orientation (Graham and Cheng, 2009a). In our experiments, most of the butterflies used the presented panorama to keep a certain heading only over a short time period and did not follow a 180° relocation of the stimulus. Apart from directional information, a panoramic skyline provides an animal with rotational optic flow information which can be used by insects for positional control (Wolf and Heisenberg, 1990). Although it is very difficult to unravel how exactly the butterflies interpreted the panorama stimulus, our data suggest that they mainly used it for flight control. Nevertheless, some individuals showed well oriented flights when presented with the panorama with high *r* values (Fig. 3A) that were not observed when the animals had optic-flow (but no distinct cue) for orientation, which indicates that these animals used the panorama for compass orientation. It will be interesting to study if these butterflies can store and memorize a desire heading with respect to the panoramic scene, a similar matching strategy to the one that has been shown in ants (Lent et al., 2010).

### Combination of multiple cues

We presented the green sun stimulus in combination with the panoramic skyline to study how the butterflies use a visual scene that mimics a combination of celestial and terrestrial information. We found that the presentation of both cues did not lead to a more directed flight performance (Fig. 4), as has been shown for the combination of multimodal cues in ants, moths, and dung beetles (Dacke et al., 2019; Dreyer et al., 2018a; Huber and Knaden, 2017; Müller and Wehner, 2007). The ability of the butterflies to keep a directed heading direction over larger time periods was dominated by the presence of the simulated sun, which is in line with observations in migrating butterflies. This suggests that the sun is the main orientation cue for monarch butterflies (Stalleicken et al., 2005). Nevertheless, we found that in the absence of the sun *or* the panorama the directedness of the butterflies was affected (Fig. 5). Some animals used both cues during flight, while other individuals relied predominantly on the simulated sun or the panoramic skyline as a reference. This indicates that the butterfly’s compass is capable of combining and weighting different visual cues, similar to what has been shown in ants and dung beetles (el Jundi et al., 2015b; el Jundi et al., 2016; Huber and Knaden, 2017). This is also similar to findings in the migrating Bogong moth which uses different modalities – the Earth’s magnetic field and dominant visual cues – for orientation (Dreyer et al., 2018a). When these cues were set in conflict, several moths were disoriented while individual moths remained well oriented. This is in line with our results that reveal a strong interindividual variability in the weighting of different orientation cues in lepidopterans and raises the question of what mechanism lepidopterans in general, and butterflies specifically, use to combine different cues in their compass. One mechanism that butterflies could use is to store multiple cues of a scene in a snapshot (with respect to the desired heading direction) and to match it to the current view, a strategy that is used by orienting dung beetles (el Jundi et al., 2016; Dacke and el Jundi, 2018). Similar to these beetles (el Jundi et al., 2015b), we know that the central complex acts as an internal compass for the butterfly’s migration (Heinze and Reppert, 2011; el Jundi et al., 2014). Thus, this brain region likely plays a major role in the integration of sun and terrestrial compass information as it provides the neuronal substrate that allows a flexible combination of different cues in the insect’s compass (Fisher et al., 2019; Kim et al., 2019; Seelig and Jayaraman, 2015). The results here show that non-migrating monarch butterflies can keep constant headings with respect to a visual scene based on skylight and terrestrial cues, similar to what migrating butterflies do during their annual journey. This suggests that the central complex controls orientation at any stage of the butterfly’s life, allowing us to study the neural mechanisms of the butterfly’s compass in detail, not only during their migration but also while they are in their non-migratory phase.

## Author contributions

Study design: MF, DD, ALS, EJW, KP, MJB, BeJ. Conducting experiments: MF, CK. Analysis of data: MF, DD, JJF, BeJ. Interpretation of data: MF, MJB, BeJ. Drafting of the manuscript: MF, BeJ. Critical review of the manuscript: CK, MJB, DD, KP, ALS, JJF, EJW. Acquired Funding: BeJ. All authors approved of the final version of the manuscript.

## Acknowledgments

We thank Konrad Öchsner for his help in designing the LED stimulus of the flight simulator. We also thank Daniel Vedder for his help in mapping the LEDs in python. We are grateful to the mechanics workshop of the Biocenter (University of Würzburg) for building important pieces of the flight simulator and Marie Dacke for her helpful comments on the manuscript. In addition, we would like to thank Sergio Siles (butterflyflyfarm.co.cr) and Marie Gerlinde Blaese for providing us with monarch butterfly pupae.

## References

Berens, P. (2009). CircStat□: A MATLAB Toolbox for Circular Statistics. J. Stat. Softw. 31, 293–295.

Brines, M. L. and Gould, J. L. (1979). Bees Have Rules. Science 206, 571–573.

Byrne, M., Dacke, M., Nordström, P., Scholtz, C. and Warrant, E. (2003). Visual cues used by ball-rolling dung beetles for orientation. J. Comp. Physiol. A 189, 411–418.

Calvert, W. H. (2001). Monarch Butterfly (*Danaus Plexippus* L., Nymphalidae) Fall Migration: Flight Behavior and Direction in Relation To Celestial and Physiographic Cues. J. Lepid. Soc. 55, 162–168.

Cartwright, B. A. and Collett, T. S. (1983). Landmark learning in bees - Experiments and models. J. Comp. Physiol. A 151, 521–543.

Coemans, M. A. J. M., Vos Hzn, J. J. and Nuboer, J. F. W. (1994). The relation between celestial colour gradients and the position of the sun, with regard to the sun compass. Vision Res. 34, 1461–1470.

Collett, T. S. and Collett, M. (2000). Path integration in insects. Curr. Opin. Neurobiol. 10, 757–762.

Collett, T. S. and Collett, M. (2002). Memory use in insect visual navigation. Nat. Rev. Neurosci. 3, 542–552.

Collett, T. S. and Land, M. F. (1975). Visual spatial memory in a hoverfly. J. Comp. Physiol. A 100, 59–84.

Dacke, M. and el Jundi, B. (2018). The Dung Beetle Compass. Curr. Biol. 28, R993–R997.

Dacke, M., Bell, A. T. A., Foster, J. J., Baird, E. J., Strube-Bloss, M. F., Byrne, M. J. and el Jundi, B. (2019). Multimodal cue integration in the dung beetle compass. Proc. Natl. Acad. Sci. U. S. A. 116, 14248–14253.

Dreyer, D., Frost, B., Mouritsen, H., Günther, A., Green, K., Whitehouse, M., Johnsen, S., Heinze, S. and Warrant, E. (2018a). The Earth’s Magnetic Field and Visual Landmarks Steer Migratory Flight Behavior in the Nocturnal Australian Bogong Moth. Curr. Biol. 28, 2160–2166.

Dreyer, D., el Jundi, B., Kishkinev, D., Suchentrunk, C., Campostrini, L., Frost, B. J., Zechmeister, T. and Warrant, E. J. (2018b). Evidence for a southward autumn migration of nocturnal noctuid moths in central Europe. J. Exp. Biol. 221, jeb179218.

Durier, V., Graham, P. and Collett, T. S. (2003). Snapshot Memories and Landmark Guidance in Wood Ants. Curr. Biol. 13, 1614–1618.

Edrich, W., Neumeyer, C. and von Helversen, O. (1979). “Anti-sun orientation” of bees with regard to a field of ultraviolet light. J. Comp. Physiol. A 134, 151–157.

el Jundi, B., Smolka, J., Baird, E., Byrne, M. J. and Dacke, M. (2014). Diurnal dung beetles use the intensity gradient and the polarization pattern of the sky for orientation. J. Exp. Biol. 217, 2422–2429.

el Jundi, B., Foster, J. J., Byrne, M. J., Baird, E. and Dacke, M. (2015a). Spectral information as an orientation cue in dung beetles. Biol. Lett. 11, 20150656.

el Jundi, B., Warrant, E. J., Byrne, M. J., Khaldy, L., Baird, E., Smolka, J. and Dacke, M. (2015b). Neural coding underlying the cue preference for celestial orientation. Proc. Natl. Acad. Sci. U. S. A. 112, 11395–400.

el Jundi, B., Foster, J. J., Khaldy, L., Byrne, M. J., Dacke, M. and Baird, E. (2016). A Snapshot-Based Mechanism for Celestial Orientation. Curr. Biol. 26, 1456–1462.

el Jundi, B., Baird, E., Byrne, M. J. and Dacke, M. (2019). The brain behind straight-line orientation in dung beetles. J. Exp. Biol. 222, 1–7.

Felt, E. P. (1925). Dispersal of Butterflies and other Insects. Nature 116, 365–368.

Fisher, Y. E., Lu, J., D’Alessandro, I. and Wilson, R. I. (2019). Sensorimotor experience remaps visual input to a heading-direction network. Nature 576, 121–125.

Fleischmann, P. N., Rössler, W. and Wehner, R. (2018). Early foraging life: spatial and temporal aspects of landmark learning in the ant *Cataglyphis noda*. J. Comp. Physiol. A 204, 579–592.

Froy, O., Gotter, A. L., Casselman, A. L. and Rappert, S. M. (2003). Illuminating the Circadian Clock in Monarch Butterfly Migration. Science. 300, 1303–1305.

Giraldo, Y. M., Leitch, K. J., Ros, I. G., Warren, T. L., Weir, P. T. and Dickinson, M. H. (2018). Sun Navigation Requires Compass Neurons in *Drosophila*. Curr. Biol. 28, 2845–2852.

Graham, P. and Cheng, K. (2009a). Which portion of the natural panorama is used for view-based navigation in the Australian desert ant? J. Comp. Physiol. A 195, 681–689.

Graham, P. and Cheng, K. (2009b). Ants use the panoramic skyline as a visual cue during navigation. Curr. Biol. 19, R935–R937.

Guerra, P. A., Merlin, C., Gegear, R. J. and Reppert, S. M. (2012). Discordant timing between antennae disrupts sun compass orientation in migratory monarch butterflies. Nat. Commun. 3:958.

Heinze, S. and Reppert, S. M. (2011). Sun compass integration of skylight cues in migratory monarch butterflies. Neuron 69, 345–358.

Heinze, S., Florman, J., Asokaraj, S., el Jundi, B. and Reppert, S. M. (2013). Anatomical basis of sun compass navigation II: the neuronal composition of the central complex of the monarch butterfly. J. Comp. Neurol. 521, 267–298.

Heinze, S., Narendra, A. and Cheung, A. (2018). Principles of Insect Path Integration. Curr. Biol. 28, R1043–R1058.

Huber, R. and Knaden, M. (2017). Homing Ants Get Confused When Nest Cues Are Also Route Cues. Curr. Biol. 27, 3706-3710.e2.

Judd, S. P. D. and Collett, T. S. (1998). Multiple stored views and landmark guidance in ants. Nature 392, 710–714.

Kim, S. S., Hermundstad, A. M., Romani, S., Abbott, L. F. and Jayaraman, V. (2019). Generation of stable heading representations in diverse visual scenes. Nature 576, 126–131.

Lebhardt, F. and Ronacher, B. (2015). Transfer of directional information between the polarization compass and the sun compass in desert ants. J. Comp. Physiol. A 201, 599–608.

Lehrer, M. and Collett, T. S. (1994). Approaching and departing bees learn different cues to the distance of a landmark. J. Comp. Physiol. A 175, 171–177.

Lent, D. D., Graham, P. and Collett, T. S. (2010). Image-matching during ant navigation occurs through saccade-like body turns controlled by learned visual features. Proc. Natl. Acad. Sci. U. S. A. 107, 16348–16353.

Merlin, C. and Liedvogel, M. (2019). The genetics and epigenetics of animal migration and orientation: Birds, butterflies and beyond. J. Exp. Biol. 222, 1–12.

Merlin, C., Gegear, R. J. and Reppert, S. M. (2009). Antennal circadian clocks coordinate sun compass orientation in migratory monarch butterflies. Science 325, 1700–1704.

Merlin, C., Heinze, S. and Reppert, S. M. (2011). Unraveling navigational strategies in migratory insects. Curr. Opin. Neurobiol. 22, 353–361.

Moore, B. R. (1980). A Modification of the Rayleigh Test for Vector Data. Biometrika 67, 175–180.

Mouritsen, H. and Frost, B. J. (2002). Virtual migration in tethered flying monarch butterflies reveals their orientation mechanisms. Proc. Natl. Acad. Sci. U. S. A. 99, 10162–10166.

Mouritsen, H., Derbyshire, R., Stalleicken, J., Mouritsen, O. O., Frost, B. J. and Norris, D. R. (2013). An experimental displacement and over 50 years of tag-recoveries show that monarch butterflies are not true navigators. Proc. Natl. Acad. Sci. 110, 7348–7353.

Müller, M. and Wehner, R. (2007). Wind and sky as compass cues in desert ant navigation. Naturwissenschaften 94, 589–594.

Narendra, A. and Ramirez-Esquivel, F. (2017). Subtle changes in the landmark panorama disrupt visual navigation in a nocturnal bull ant. Philos. Trans. R. Soc. B Biol. Sci. 372: 20160068.

Narendra, A., Gourmaud, S. and Zeil, J. (2013). Mapping the navigational knowledge of individually foraging ants, *Myrmecia croslandi*. Proc. R. Soc. B 280: 20130683.

Philippides, A., Baddeley, B., Cheng, K. and Graham, P. (2011). How might ants use panoramic views for route navigation? J. Exp. Biol. 214, 445–451.

Reid, S. F., Narendra, A., Hemmi, J. M. and Zeil, J. (2011). Polarised skylight and the landmark panorama provide night-active bull ants with compass information during route following. J. Exp. Biol. 214, 363–370.

Reppert, S. M. (2006). A colorful model of the circadian clock. Cell 124, 233–236.

Reppert, S. M. and de Roode, J. C. (2018). Demystifying Monarch Butterfly Migration. Curr. Biol. 28, R1009–R1022.

Reppert, S. M., Zhu, H. and White, R. H. (2004). Polarized Light Helps Monarch Butterflies Navigate. Curr. Biol. 14, 155–158.

Reppert, S. M., Guerra, P. A. and Merlin, C. (2016). Neurobiology of Monarch Butterfly Migration. Annu. Rev. Entomol. 61, 25–42.

Rossel, S. and Wehner, R. (1984). Celestial orientation in bees: the use of spectral cues. J. Comp. Physiol. A 155, 605–613.

Sauman, I., Briscoe, A. D., Zhu, H., Shi, D., Froy, O., Stalleicken, J., Yuan, Q., Casselman, A. and Reppert, S. M. (2005). Connecting the navigational clock to sun compass input in monarch butterfly brain. Neuron 46, 457–467.

Seelig, J. D. and Jayaraman, V. (2015). Neural dynamics for landmark orientation and angular path integration. Nature 521, 186–191.

Stalleicken, J., Mukhida, M., Labhart, T., Wehner, R., Frost, B. and Mouritsen, H. (2005). Do monarch butterflies use polarized skylight for migratory orientation? J. Exp. Biol. 208, 2399–2408.

Stevens, V. M., Turlure, C. and Baguette, M. (2010). A meta-analysis of dispersal in butterflies. Biol. Rev. 85, 625–642.

Warrant, E., Frost, B., Green, K., Mouritsen, H., Dreyer, D., Adden, A., Brauburger, K. and Heinze, S. (2016). The australian bogong moth *Agrotis infusa*: A long-distance nocturnal navigator. Front. Behav. Neurosci. 10, 1–17.

Wehner, R. (1997). The ant’s celestial compass system: spectral and polarization channels. In Orientation and Communication in Arthropods (ed. Lehrer, M.), pp. 145–185. Basel: Birkhäuser.

Wehner, R. and Müller, M. (2006). The significance of direct sunlight and polarized skylight in the ant’s celestial system of navigation. Proc. Natl. Acad. Sci. U. S. A. 103, 12575–12579.

Wolf, R. and Heisenberg, M. (1990). Visual control of straight flight in Drosophila melanogaster. J. Comp. Physiol. A 167, 589–592.

